# Progressive postnatal hearing development limits early parent-offspring vocal communication in the zebra finch

**DOI:** 10.1101/2024.05.03.592470

**Authors:** Tommi Anttonen, Jakob Christensen-Dalsgaard, Coen P. H. Elemans

## Abstract

Acoustic communication relies critically on the receiver’s ability to hear. In precocial birds, hearing can already be functional during embryonic stages in the egg, whereas in altricial species the limited available data indicate a gradual postnatal development with poor sensitivity in early hatchlings, implying even lower sensitivity in embryos. Recent research suggests that zebra finch embryos, despite being altricial, have functional hearing already in the egg to engage in parent-embryo acoustic communication and anthropogenic noise detection. However, their auditory sensitivity during early development remains unknown. Here, we measure auditory brainstem responses (ABR) over early postnatal development and show that zebra finch hatchlings have >54 dB lower threshold than adults to loud, broadband clicks. ABR responses emerge between 4-8 days after hatching. Auditory sensitivity develops progressively and reaches adult levels by day 20. Additionally, egg vibrations induced by sound remain far below detection thresholds of vibrotactile senses. Together these data contradict the notion of acoustic perception in embryos and early parent-to-embryo communication in zebra finches. The striking timing coincidence between maturation of the peripheral auditory system and the onset of song learning suggests that hearing functionality may gate the onset of vocal learning in zebra finches.

## Introduction

The ability to detect sound is critical for vocal communication, species recognition, predator detection, and survival across the vertebrates (Nordeen and Nordeen, 1992; Heaton et al., 1999; Zevin et al., 2004; Ladich and Winkler, 2017). In birds, precocial species that are able to move around after hatching show some hearing function already inside the egg (Gottlieb, 1965; Saunders et al., 1973; Konishi, 1973). This enables embryo-to-embryo (Vince, 1966; Woolf et al., 1976; Schwagmeyer et al.,1991; Noguera and Velando, 2019) and parent-to-embryo communication that is critical to imprinting on species-specific sounds (Gottlieb, 1965; Grier et al., 1967; Hess, 1972). In contrast, altricial birds hatch considerably earlier in their development and rely on parental care to survive (Orkney et al. 2025). The sparse data available show indicate a gradual postnatal development of hearing sensitivity, with poor sensitivity in early hatchlings (Brittan-Powell and Dooling, 2004; Korneeva et al., 2006; Aleksandrov and Dmitrieva, 1992; Kraemer et al., 2017), implying even lower sensitivity in the less developed prenatal embryos.

Remarkably, recent research directly challenges this concept and suggests that altricial songbirds can detect soft sounds while still in the egg (from here onwards referred to as embryos). In zebra finches, embryonic exposure to high-frequency “heat-call” (from here onwards referred to as heat whistles (Anttonen et al., 2025) playbacks has been associated with differences in physiological and behavioral traits (Mariette and Buchanan, 2016; Mariette et al., 2018; Katsis et al., 2018; Mariette and Buchanan, 2019; Pessato et al., 2020; Udino et al., 2021; Katsis et al., 2021; Udino and Mariette, 2022; Katsis et al., 2023) and prehatching anthropogenic noise exposures reduced embryonic survival and reproductive success in adulthood (Meillère et al., 2024). Furthermore, fairy wren embryos supposedly learn elements of the female incubation call already at 10-12 days after fertilization (Colombelli-Négrel et al., 2012, 2014, 2016; Kleindorfer et al., 2024). The above interpretations critically depend on the ability of these embryos to detect soft level sounds in the egg and after hatching. However, it is currently unknown whether these species can detect sounds this early in their development.

The zebra finch is a major animal model system for studying the neural circuitry underlying hearing-guided behaviors including vocal imitation learning. However, we know surprisingly little about the development and the function of the auditory organ in this species. In singing males, the onset of sensory and sensorimotor learning starts around 25 days-post-hatch (DPH) and continues to adulthood at ∼90 DPH (Immelmann, 1967; Böhner, 1990; Roper and Zann, 2006; Braaten, 2010; Gobes et al., 2019; Leitão and Gahr, 2024). Aligned with the onset of this behavior, the overall hearing sensitivity is comparable to adult levels at 20 DPH (Amin et al., 2007) with a frequency range from 0.25 to 8 kHz and highest sensitivity around 1-6 kHz (Behavior 1-6 kHz: Okanoya and Dooling, 1987; Yeh et al., 2023; ABR 1-4 kHz, Amin et al., 2007; Noirot et al., 2011; Zevin et al 2004). However, at 10 DPH - the earliest age measured – hearing sensitivity is reduced by 25 dB compared to 20 DPH and is thus ∼20-fold lower (Amin et al., 2007). Because hearing sensitivity in zebra finches increases progressively in late postnatal development - as in other bird species (Rebillard and Rubel, 1981; Katayama, 1985; Jones et al., 2006; Brittan-Powell and Dooling, 2004; Köppl and Nickel, 2007; Kraemer et al., 2017) their ability to detect sounds as embryos and as early hatchlings may be severely compromised. Furthermore, recent work shows that heat whistles are extremely soft sounds *in vivo* with a source level of only ∼14 dB re. 20 µPa at 1 meter (sound pressure level of ∼34 dB re. 20 µPa at 10 cm; Anttonen et al., 2025). Therefore, zebra finch embryos would require highly sensitive hearing function to detect such soft sounds. However, we lack conclusive evidence on the hearing capabilities of juvenile zebra finches.

Here we test the hypothesis that zebra fnches can hear during early postnatal development by measuring auditory brainstem responses (ABRs) that allow hearing threshold estimation when behavioral training is not possible (He et al., 2008). Furthermore, we test whether intense sounds can induce egg vibrations of sufficient magnitude for vibrotactile detection. We show that the earliest ABRs evoked by intense (95 dB), broadband sounds could not be detected on 2 DPH, but earliest at 4-8 DPH. The threshold at 4DPH is at least >54 dB higher compared to adult levels. Responses to frequency specific sounds develop later in a progressive, low-to-high frequency order. ABR sensitivity to clicks matures to adult-like levels at 10 DPH but the response amplitude continues to increase beyond 20 DPH. Airborne sounds induce low amplitude vibrations in eggs that are far below typical avian and mammalian vibrotactile detection thresholds. Together, our findings show that zebra finch embryos and early hatchlings are functionally deaf and provide evidence against the hypothesis that they can detect sound stimuli via hearing or via vibrotactile senses. Our data furthermore show that auditory nerve responses continue to mature past 20 DPH, suggesting that the maturation of hearing limits the onset of vocal learning in zebra finches.

## Results

### Responses to loud clicks appear during the first postnatal week

To assess the development of zebra finch hearing sensitivity in early postnatal development we quantified the responses of auditory pathway neurons to sound stimuli by recording ABRs (**Fig. 1A**). First, to evaluate overall hearing sensitivity we presented loud yet physiologically relevant (95 dB SPL) clicks that stimulate the auditory system across a wide frequency spectrum (**Fig. 1B**). Clicks activate most auditory neurons nearly simultaneously and result in prominent ABRs, thus providing maximal sensitivity for threshold detection.

**Figure 1.**
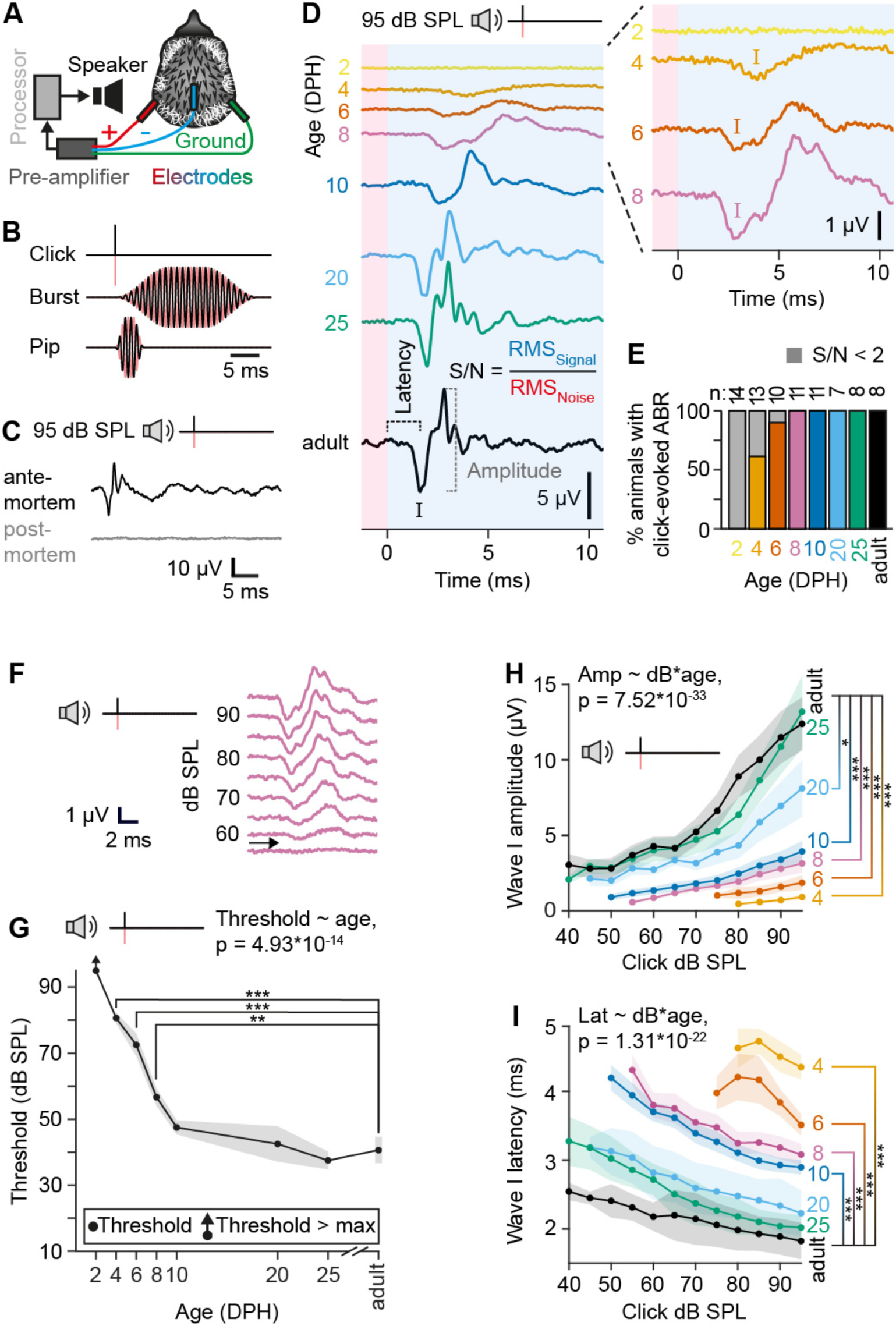
Click-evoked ABRs appear and gradually mature after hatching in zebra finches. (**A**) ABR recording setup. (**B**) Used sound stimuli. *Red*, phase inverted signal. (**C**) Representative, 95 dB SPL click-evoked ABR *ante-*(*black*) and *post-mortem* (*grey*). (**D**) Representative ABRs to 95 dB SPL clicks at tested DPH timepoints. The first negative peak (*I*) and the following trough are designated as wave I. Noise RMS is determined from the *red area* and signal RMS from the *blue area*. (**E**) Percentage of animals that exhibited a detectable (S/N > 2) click-evoked ABR. (**F**) Detection thresholds were measured with a set of clicks with decreasing 5 dB SPL steps. *Arrow* indicates the determined threshold for this 8 DPH example. (**G**) Average click thresholds at different DPH ages. (**H**) Click-evoked ABR wave I amplitude and (**I**) latency over SPL. Age groups are *color-coded* in D-I. *Shaded areas* in H-I = s.e.m.

Clicks induced robust neural potentials in adult zebra finches (**Fig. 1D-E**). These potentials were not recording artefacts as they occurred only when the animal was alive (N = 3, **Fig. 1C**). None of the tested animals displayed click-evoked ABRs at 2 DPH (both with detection at signal-to-noise ratio of 2 (S/N > 2) and with detailed visual inspection; n=0/15; **Fig. 1D,E**). However, click-evoked ABRs became observable in most animals during the first postnatal week (S/N > 2; 4 DPH, n = 8/13; 6 DPH, n = 9/10; **Fig. 1D,E**). Subsequently, all tested animals consistently demonstrated click-evoked ABRs from the second postnatal week onwards (S/N > 2; 8 DPH, n = 11; 10 DPH, n = 11; 20 DPH, n = 7; 25 DPH, n = 8; Adult, n = 8; **Fig. 1D,E**). Thus, sounds at loud, yet physiologically relevant SPLs do not evoke ABRs in the first days after hatching, but do so in all animals at 8 DPH.

### Click-evoked ABR thresholds are adult-like at 10 DPH

To estimate hearing sensitivity thresholds over postnatal age, we recorded series of click-evoked ABRs at 5 dB downward steps from 95 dB SPL (**Fig. 1F**). Hearing thresholds are strongly affected by age of the animal (F(6) = 25.22, p = 4.93*10^-14^; **Fig. 1G**, **Table S1**). Click-evoked ABR thresholds are significantly higher than adult-like levels (40.6 ± 4.0 dB SPL) in animals younger than 10 DPH but not in older animals (4 DPH vs. adult, t = 8.16, p = 3.27*10^-10^; 6 DPH vs. adult, t = 5.92, p = 1.39*10^-6^; DPH 8 vs. adult, t = 3.37, p = 0.008; 10 DPH vs. adult, t = 1.23, p = 1; 20 DPH vs. adult, t = -0.13, p = 1; 25 DPH vs. adult, t = -1.29, p = 1; p-values are Bonferroni-corrected; **Fig. 1G**, **Table S1**). Click thresholds are above 95 dB on 2 DPH and as high as 80.6 ± 1.6 dB SPL on 4 DPH. The threshold of a 2 DPH animal is at least 54 dB higher compared to an adult. Thus, zebra finch hearing sensitivity increases rapidly but gradually during the first two postnatal weeks.

### ABR wave I gradually reaches maturity 25 days after hatching

The click-evoked ABR consists of multiple wave peaks and troughs that reflect the subsequent activations of various neural populations of the auditory pathway. The first wave (wave I in **Fig. 1D**) is considered to correspond to the activity of the primary auditory neurons (Brittan-Powell and Dooling 2002). To study the postnatal maturation of this synchronized neural output, we measured changes in both the amplitude and the latency of wave I over postnatal development.

In adults, the amplitude of wave I increases significantly with click SPL (**Fig. 1H**). Age significantly affects both wave I amplitude directly and the relation between SPL and wave I amplitude (Age-SPL interaction compared to only additive effects of age and SPL: *𝜒*^2^(6)=164.23, p=7.52*10^-33^, additive effects of age and SPL compared to only SPL: *𝜒*^2^(6)=79.43, p=4.69*10^-15^, SPL only compared to null model: *𝜒*^2^(1)=438.12, p=2.78*10^-97^, **Fig. 1H**, **Table S2**). The steepness of the amplitude-SPL slope gradually increases with age and is significantly different between adults and younger animals, but not between adult and those aged 25 DPH (4 DPH vs. adult, t = -5.40, p = 1.50*10^-6^; 6 DPH vs. adult, t = -5.65, p = 1.8910^-6^; DPH 8 vs. adult, t = -6.30, p = 3.79*10^-7^; 10 DPH vs. adult, t = -5.53, p = 118 6.53*10^-6^; 20 DPH vs. adult, t = -3.00, p = 0.03; 25 DPH vs. adult, t = -0.69, p = 1; p-values are Bonferroni-corrected; **Fig. 1H**, **Table S2**).

The latency of wave I decreases with increasing click SPL (**Fig. 1I**). Furthermore, age significantly affects the relation between dB SPL and wave I latency (Age-SPL interaction compared to only additive effects of age and SPL: *𝜒*^2^(6) = 116, p = 1.31*10^-22^; additive effects of age and SPL compared to only SPL, *𝜒*^2^(6) = 109, p = 2.76*10^-21^; SPL only compared to null model: *𝜒*^2^(1) = 754, p = 4.81*10^-166^; **Fig. 1I**, **Table S3**). The observed latencies are significantly longer during early postnatal development (4-to-10 DPH) when compared to latencies observed in adults. No significant differences are observed between the latencies of adults and animals older than 10 DPH (4 DPH vs. adult, t = 12.56, p = 3.41*10^-19^; 6 DPH vs. adult, t = 10.06, p = 1.49*10^-13^; 8 DPH vs. adult, t = 7.24, p = 1.16*10^-8^; 10 128 DPH vs. adult, t = 6.18, p = 5.95*10^-7^; 20 DPH vs. adult, t = 2.24, p = 0.18; 25 DPH vs. adult, t = 1.67, 129 p = 0.61; p-values are Bonferroni-corrected; **Fig. 1I**, **Table S3**).

Taken together, the synchronized output of the auditory nerve increases in sensitivity and reduces in latency postnatally progressively from 4 DPH until reaching maturity at 20-25 DPH. Such gradual increase is consistent with postnatal hearing development in budgerigars, mice, and humans (Brittan-Powell and Dooling, 2002, 2004; Hecox and Galambos, 1974; Henry and Haythorn, 1978).

### Hearing sensitivity develops progressively from low to high frequencies

To study the postnatal development of frequency-specific hearing sensitivity we tested whether loud (95 dB SPL) tone bursts centered at frequencies between 0.25-to-8 kHz elicit detectable ABRs. Tone burst-evoked ABRs were first detectable on 8 DPH at 1 kHz (**Fig. 2A** and **Fig. S1A**). As observed with click stimuli, the amplitude of tone burst-induced ABRs increases with age (**Fig. 2A**, **Fig. S1B, Table S4**; Age as a predictor of amplitude compared to null model: *𝜒*^2^(3) = 6.11, p = 0.11). The latency of the response decreases significantly with age (**Fig. 2a**, **Fig. S1c, Table S4**; Age as predictor of latency compared to null model: *𝜒*^2^(3) = 33.72, p = 2.27*10^-7^). Latencies are significantly longer at 8-to-10 DPH than at adulthood (1 kHz: 8 DPH, 6.60 ± 0.77 ms; 10 DPH, 5.01 ± 0.40 ms) and are adult-like at 20-to-25 DPH (1 kHz: 20-to-25 DPH, 3.34 ± 0.18 ms; adult, 3.67± 0.30 ms; 8 DPH vs. adult, t = 5.95, p = 1.87*10^-6^; 10 DPH vs. adult, t = 4.52, p = 0.0004; 20-to-25 DPH vs. adult, t = 1.16, p = 0.81; p-values are Bonferroni-corrected; **Fig. S1C** and **Table S4**). During the following weeks of postnatal development, tone burst-induced ABRs became detectable at 0.5-to-4 kHz (**Fig. S1A**). No responses were detected at higher frequencies (**Fig. 2B,C**; **Fig. S1A**). Taken together, the maturation of hearing sensitivity in zebra finches progresses in a low-to-high frequency order as is also observed for example in mice (Ehret, 1976), cats (Ehret and Romand, 1981), and in some other bird species (Brittan-Powell and Dooling, 2002, 2004; Köppl and Nickel, 2007; Kraemer et al., 2017).

**Figure 2.**
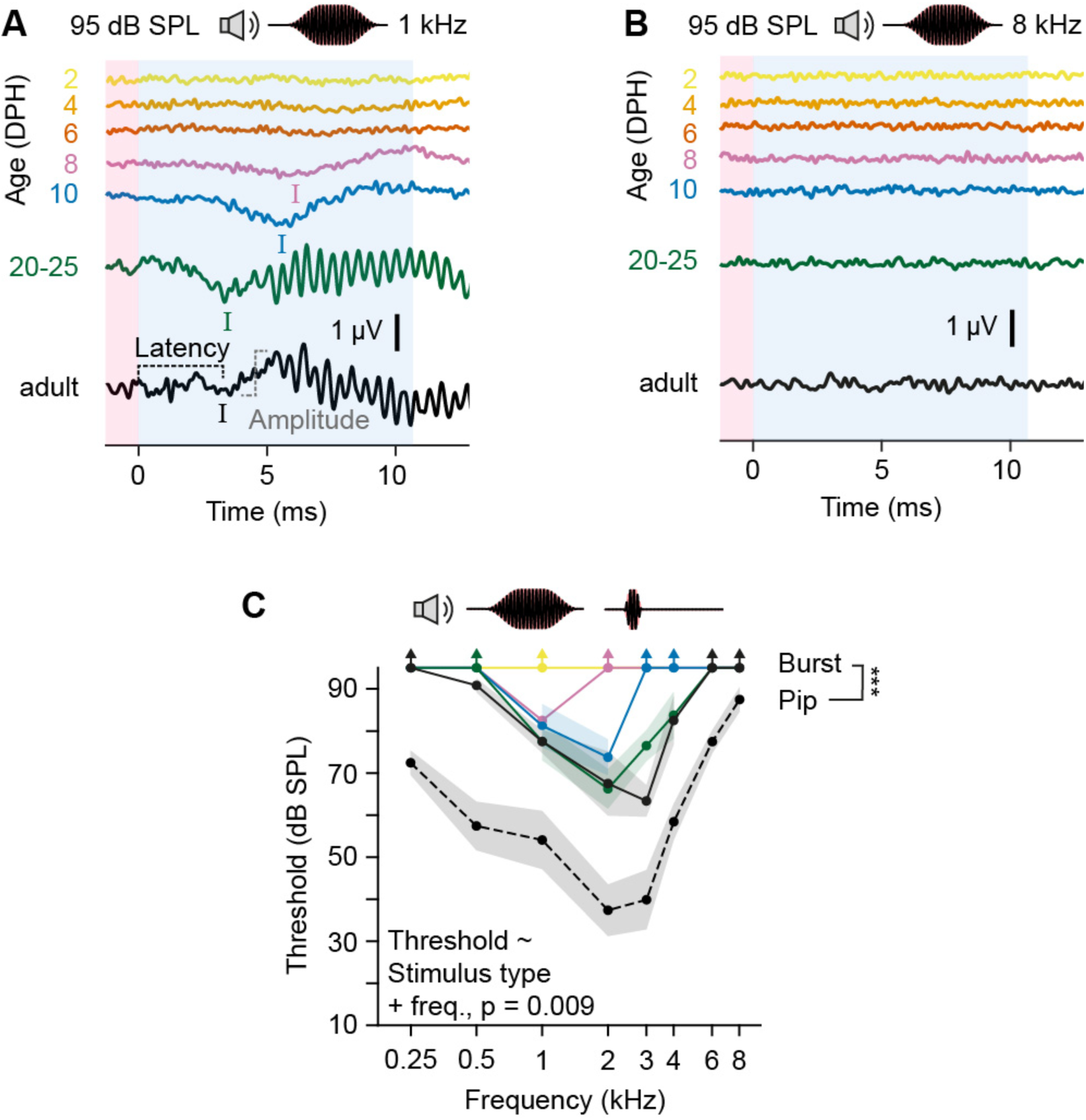
Sound frequency sensitivity matures over postnatal development. (**A**) Representative ABRs to 1 kHz and (**B**) 8 kHz tone burst stimuli over postnatal development. (**C**) Zebra finch ABR thresholds obtained with tone bursts (*solid lines*) and with tone pips (*dashed line*).) *Shaded areas* = s.e.m.

### Tone pip-evoked ABR thresholds

Tone bursts evoked ABRs only in about half of the adults (**Fig. S1A**) and the detection thresholds of > 60 dB SPL are high compared to earlier ABR studies (Amin et al., 2007; Noirot et al., 2011; Zevin et al 2004). This indicates that while being more frequency specific, tone bursts failed to recruit auditory neurons to fire concurrently enough to generate detectable ABRs. Therefore, we next used shorter 5 ms-long tone pips (**Fig. 1B**) that have a broader spectral distribution of acoustic energy compared to tone bursts, but produce a more focused temporal response (Lauridsen et al., 2021).

Indeed, the SPL matched tone pips produced much more prominent ABRs (**Fig. 2C** and **Fig. S2A**) that were readily detected in all tested adults around their most sensitive hearing range of 1-to-4 kHz (**Fig. S2B**). Detection thresholds for tone pips were significantly lower than those of tone bursts with ∼20 dB SPL (Additive effects of stimulus type and frequency as predictors of threshold compared to stimulus type alone χ^2^(1) = 6.84, p = 0.009; stimulus type as a predictor of threshold compared to null model: χ^2^(1) = 10.27, p = 0.001, **Table S5**).

### Sound-induced egg vibrations

These data suggest it unlikely that embryonic zebra finches can detect soft, high frequency heat whistles as airborne sounds via their auditory system (See Discussion). Therefore, we next tested if airborne sounds can induce zebra finch eggs to move at sufficient velocities to be detected as vibrations.

We played sound frequency sweeps to eggs while simultaneously recording the sound-induced vibrations of the egg with a laser vibrometer (**Fig. 3**). The vibration velocity transfer function shows that eggs vibrate at velocities around -85 to -55 dB re 1 mm s^-1^ when exposed to a loud 94 dB SPL frequency sweep between 0.25-to-10 kHz (**Fig. 3C**). Within the frequency ranges associated with zebra finch song (1-4 kHz, Elie and Theunissen 2016) and heat whistles (6.8 ± 0.6 kHz (Anttonen et al, 2025), sound exposures at 94 dB SPL induce average vibration velocities of -72.4 ± 0.8 and -70.7 ± 1.2 dB re 1 mm s^-1^(N=5), respectively (**Fig. 3B**). We calculated the corresponding vibration velocity levels induced by SPLs used during experimental heat whistle playbacks to zebra finch eggs (62-to-67 dB SPL)(Katsis et al., 2018) to be between -99.4 and -96.2 dB re 1 mm s^-1^, respectively (**Fig. 4**).

**Figure 3.**
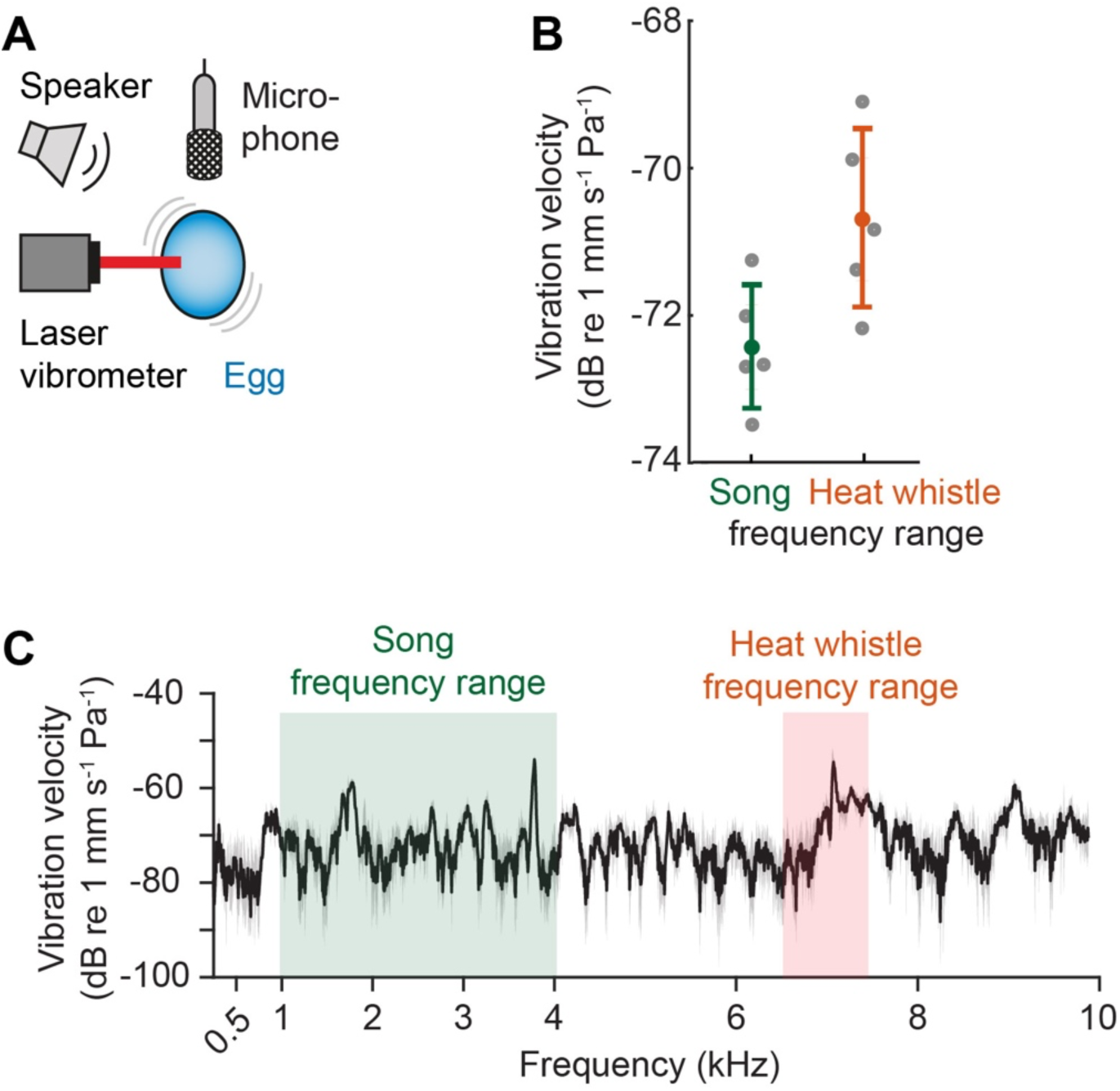
Intense sound stimulation induces zebra finch eggs to vibrate at very low velocities. **A** Setup used to measure sound induced vibrations of eggs. A 94 dB, 0.25-to-10 kHz frequency sweep was played at the eggs to determine the vibration transfer function. **B** Egg vibration velocities at frequencies associated with song (*left*) and with heat whistles (*right*). Error bars = s.d. **C** Example vibration velocity transfer function of a zebra finch egg (mean ± S.D. (grey area) N=5) *Green* and *red areas* display the frequency ranges of song (Elie and Theunissen 2016) and heat whistles (Anttonen et al, 2025), respectively, that were used to calculate mean velocity in panel B.

**Figure 4.**
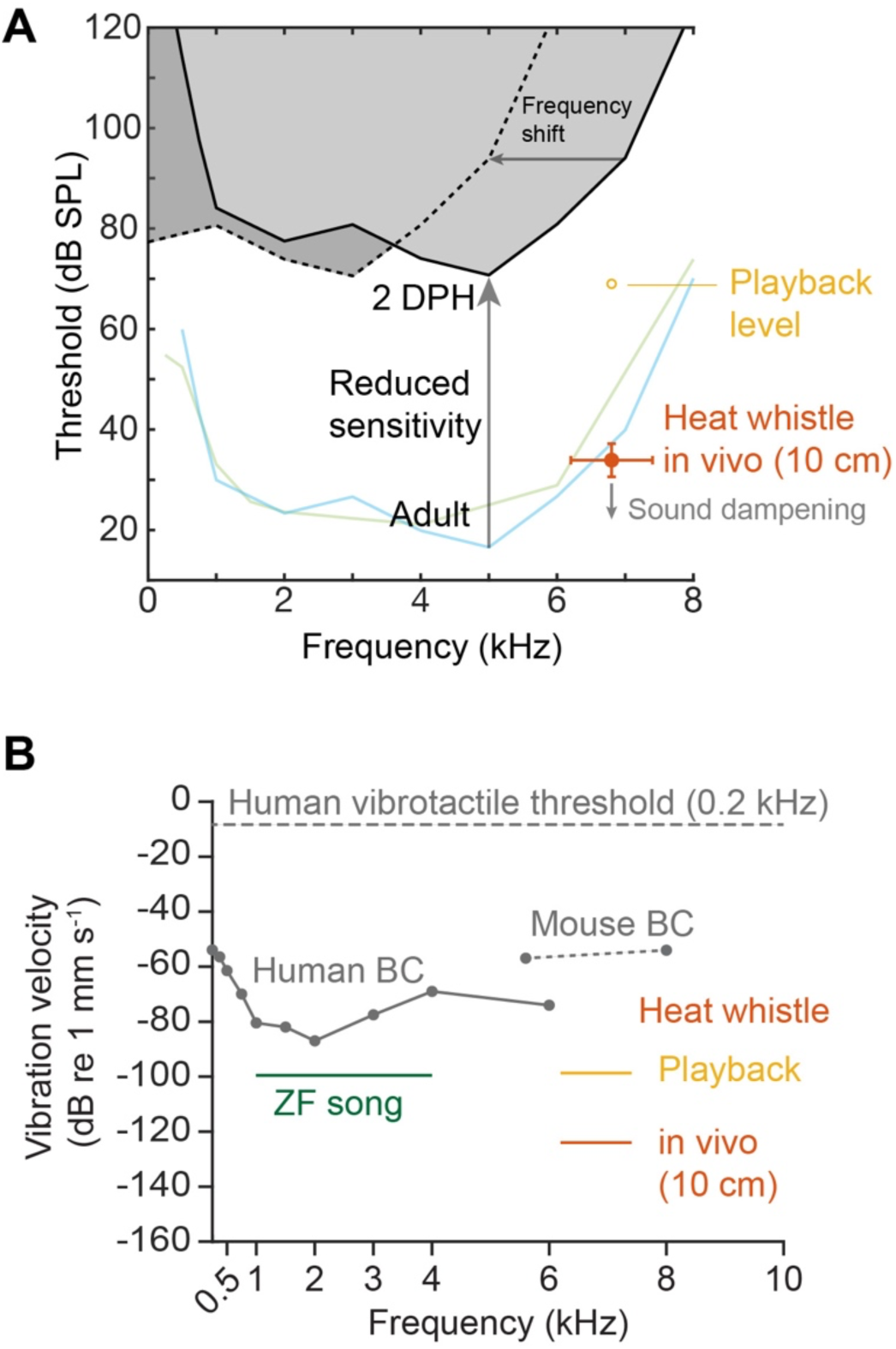
Heat whistles are below hearing and vibration threshold levels in hatchlings and adults. **A** *In vivo* heat whistles at 10 cm distance (Anttonen et al., 2025) (*orange*) are just on or below the adult hearing threshold based on behavioral audiograms (blue line; Yeh et al. 2023; green line, Okanoya and Dooling, 1987). Increasing the most sensitive adult audiogram by 54 dB as a conservative estimate of reduction based in clicks (Fig 1G), shows that both in vivo heat whistles and previously playback levels (Katsis et al., 2018) lay far below the estimated thresholds for a 2DPH old hatchling, even without an expected further increased high frequency threshold (horizontal arrow). Thresholds for embyos still in the egg are likely be shifted further up, and sound dampening by parents incubating the egg will reduce the heat whistle levels further down, making the gap even larger. **B** Egg vibration velocities caused by 67 dB song, 65 dB heat whistle playbacks (Katsis et al., 2018), and in vivo heat whistles (Anttonen et al., 2025) are far below with thresholds for human (Håkansson et al., 1985) and mouse bone conduction (BC) hearing (Chhan et al., 2017) and for human vibrotactile threshold at 0.2 kHz (Mountcastle et al., 1972).

## Discussion

We show that hearing functionality gradually increases over postnatal development in zebra finches. An evoked response (ABR) to loud click stimuli, reflecting the synchronized output of auditory nerve fibers, is first detectable at 4-8 days post hatch (DPH) after which hearing sensitivity gradually increases until reaching adult levels at 20-25 DPH. The response to longer tone burst also gradually increases consistent with hearing development in altricial and precocial bird species (Saunders et al., 1973; Konishi, 1973; Rebillard and Rubel, 1981; Katayama, 1985; Jones et al., 2006; Brittan-Powell and Dooling, 2004; Köppl and Nickel, 2007; Kraemer et al., 2017; Aleksandrov and Dmitrieva, 1992; Lippe, 1994; Amin et al., 2007) - as well as in mammals (Henry and Haythorn, 1978; Ehret and Romand, 1981). Therefore, the functional maturation of hearing sensitivity over postnatal development in zebra finches follows a gradual, progressive pattern consistent with other birds and mammals.

The progressively increasing hearing sensitivity over postnatal development (**Fig. 1**) implies that zebra finch embryos have a further reduced hearing sensitivity compared to early hatchlings. Even in precocial bird species that show auditory function already in the egg, auditory thresholds remain very high until a few days before hatching (Saunders et al., 1973; Rebillard and Rubel, 1981; Lippe, 1994; Saunders et al., 1974). In precocial chickens, evoked responses to very loud sounds (>100 dB SPL) have been obtained around embryonic day 12 (chickens hatch at day 21) (Saunders et al., 1973), but ambient sound levels evoke responses around four days later (Jones et al., 2006). Importantly, these measurements involved experimentally opening the egg, clearing the auditory meatus from fluids, and even drainage of fluids from the middle ear (Rebillard and Rubel, 1981) and thus may overestimate the natural hearing sensitivity. Although ABR provides easily accessible measurements of hearing sensitivity, the technique does not inform on the anatomical and neural processes underlying the gradual development of hearing sensitivity. In birds, these processes have been studied in most detail in chickens, where fluid unloading and maturation of middle ear structures causes significant improvements in hearing sensitivity during development (Saunders et al., 1973; Cohen et al., 1992). The onset of evoked responses to very loud sounds furthermore coincides with synapse formation on hair cells of the inner ear (Saunders et al., 1973; Cohen and Fermin, 1978; Caus Capdevila et al., 2021). Furthermore, the response characteristics of the auditory nerve fibers and the latencies of the ABR show postnatal changes (Katayama,1985; Jones et al., 2006; Manley et al., 1991). The periphery of the chicken auditory system also continues to anatomically mature for several weeks after hatching (Cohen et al., 1992; Ryals et al., 1984; Cotanche and Corwin, 1991). Obviously, precocial species hatch much later in their development than altricial species and when these hallmark events occur in zebra finch development remains unknown. Further work is required to understand the physiological basis for postnatal development of hearing sensitivity in the zebra finch and other songbird species.

Our data indicate that zebra finch embryos and early hatchlings are functionally deaf within the tested stimulus range. ABR measures synchronized population activity and therefore provides a conservative estimate of sensitivity (Pickles, 2012). In clinical practice, the absence of ABR responses at high intensities is used to diagnose severe or profound hearing impairment in human infants (Stapells, 2000; WHO, 2021), with functional deafness typically defined at thresholds ≥81 dB. By this standard, the lack of detectable responses at the highest stimulus levels tested (95 dB SPL) reflects a profound reduction in functional sensitivity. While sparse activity below ABR detection thresholds cannot be excluded, such activity is unlikely to support meaningful perception or behavior. We therefore use “functional deafness” to denote the absence of detectable, population-level neural responses under physiologically relevant conditions, rather than absolute absence of any neural activity.

When in a hot environment, zebra finch parents emit heat whistles that are soft, high frequency sounds (∼7 kHz, Anttonen et al., 2025; 6-10 kHz, Katsis et al., 2018) compared to zebra finch song, that are produced during the high respiratory flow conditions consistent with thermal panting (Anttonen et al., 2025; McDiarmid et al., 2018; Pessato et al., 2020). Embryonic exposure to heat whistle playbacks has been associated with differences in physiological and behavioral traits (Mariette and Buchanan, 2016; Mariette et al., 2018; Katsis et al., 2018; Mariette and Buchanan, 2019; Pessato et al., 2020; Udino et al., 2021; Katsis et al., 2021; Udino and Mariette, 2022; Katsis et al., 2023). Our data provide no support for the hypothesis that zebra finch embryos or early hatchlings can detect heat whistles via either auditory or vibrotactile pathways (**Fig 4**). This conclusion does not rely on a single dataset or method, but on the convergence of three independent constraints: (i) heat whistle sound level, (ii) adult auditory sensitivity, and (iii) developmental immaturity of the auditory system (**Fig 4a**). First, heat whistles are low-amplitude sounds. In vivo measurements show sound levels of ∼33 dB SPL at 10 cm and ∼14 dB at 1 m, with a conservative upper bound of ∼40 dB SPL even at very close range (<5 cm) (Anttonen et al., 2025). Second, behavioral audiograms indicate that adult zebra finches have thresholds of ∼30 dB SPL around 7 kHz, increasing steeply at higher frequencies (Okanoya and Dooling, 1987; Yeh et al., 2023) (**Figs 4, 5**). Thus, even under optimal conditions and at close range (10 cm), heat whistles lie at or below the detection threshold of adults (**Fig 4A**). Third, auditory sensitivity in early development is substantially reduced: our ABR data show a ≥54 dB loss in sensitivity in hatchlings relative to adults for click stimuli. In addition, the progressive frequency-dependent development from low to high suggests that the high frequency (6–8 kHz) thresholds can be expected to be even higher (left frequency shift in Fig **4A**). Together, these constraints define a narrow and unfavorable detection window—a low-amplitude, high-frequency sound positioned at the edge of adult sensitivity, combined with a large developmental reduction in sensitivity—making detection by embryos or hatchlings highly unlikely. Importantly, this conclusion is robust to uncertainties in ABR–behavioral offsets; even under the most conservative assumptions, the sound remains at or below adult detection limits and far below those expected in early development.

Consistent with this, comparative data show that high-frequency hearing matures late across vertebrates, and even precocial species with embryonic auditory function exhibit very poor sensitivity above 6 kHz prior to hatching (Saunders et al., 1973; Katayama, 1985). Established parent-embryo communication in precocial mallard ducks depends specifically on the auditory sensitivity of the embryo to detect maternal calls with frequency components below 3 kHz (Gottlieb, 1975). We further rule out a vibrotactile or bone-conduction pathway (**Fig 4B**): sound-induced egg vibrations by heat whistle playbacks (65 db SPL (Katsis et al., 2018)) are 10–40 dB below vertebrate bone-conduction thresholds and ∼90 dB below vibrotactile detection thresholds (Håkansson et al., 1985 (human bone conduction); Mountcastle et al., 1972 (human and monkey vibrotactile); Hörster, 1990 (pigeon vibrotactile)). *In vivo* heat whistles are at least 27 dB weaker than the used playback levels (Anttonen et al., 2025). Even assuming lossless transmission to the embryo, these vibrations remain orders of magnitude below known detection limits. Moreover, parental incubation would attenuate airborne sound while generating substantially stronger mechanical cues through respiration and contact, which were not the signals proposed to mediate these effects. Taken together, both auditory and mechanical pathways fall well below physiological detection limits, and are not plausible mechanism by which heat whistles could be perceived by embryos under natural conditions.

Taken together, our results provide evidence against the hypothesis that zebra finch embryos can hear sounds, and we consider it thus unlikely that adult zebra finches can communicate with their embryos via air-borne sound. How the heat whistle and anthropogenic noise playbacks have affected the embryos in the earlier studies (Mariette and Buchanan, 2016; Mariette et al., 2018; Katsis et al., 2018; Mariette and Buchanan, 2019; Pessato et al., 2020; Udino et al., 2021; Katsis et al.,2021; Udino and Mariette, 2022; Katsis et al., 2023; Meillère et al., 2024) remains to be further tested. Resolving this issue will require experiments that integrate realistic in vivo sound levels with simultaneous measurement of auditory, vibrotactile, and thermal cues. Playback studies should be calibrated to natural sound amplitudes (Anttonen et al., 2025), and complemented by approaches with higher sensitivity than ABR, such as single-unit recordings or optical imaging. A comparative framework spanning altricial and precocial species, combined with developmental staging based on anatomical and neural maturation of the auditory system and novel behavioral essays, will be essential to define the limits of sensory function in the avian embryo.

Behavioral studies show that altricial songbird nestlings suppress begging in response to parental alarm calls and predator cues several days after hatching, typically emerging between ∼5–10 days post-hatch and strengthening thereafter (Platzen and Magrath, 2004; Magrath et al., 2006; Magrath et al., 2007; Haff and Magrath, 2012; Barati and McDonald, 2017; Suzuki, 2011). This timeline closely matches the onset and maturation of ABR responses observed here in zebra finches. Importantly, these behaviors likely rely on detection of salient, often low-frequency or broadband cues, and do not imply mature auditory sensitivity. Korneeva et al. (2006) and Aleksandrov and Dmitrieva (1992) report ABR responses in 3DPH flycatcher hatchlings using invasive recordings in unanesthetized birds, which likely yield lower thresholds. Even so, sensitivity improves rapidly postnatally, with ∼40 dB threshold decreases and an upper frequency shift from ∼4 to ∼7 kHz within days. Our physiological data therefore align with the literature, indicating that ecologically relevant auditory function emerges only after an initial postnatal period of markedly reduced sensitivity.

Our adult pip audiograms are broadly consistent with values reported across multiple laboratories (**Fig. 5**), despite substantial methodological variation (Amin et al., 2007; Noirot et al., 2011; Zevin et al 2004). Differences in electrode configuration (subdermal vs. in-brain), stimulus duration and envelope, repetition rate, and threshold criteria are all known to influence ABR thresholds and amplitudes (Stapells, 2000; Brittan-Powell et al., 2002), yet the results from four laboratories with different setups, conditions and bird populations are highly comparable. The remaining spread across studies likely reflects both technical and biological sources, including colony-specific variation and the limited representation of wild-derived zebra finches. Standardized protocols across populations will be required to disentangle methodological from biological variability in avian ABR datasets.

**Figure 5.**
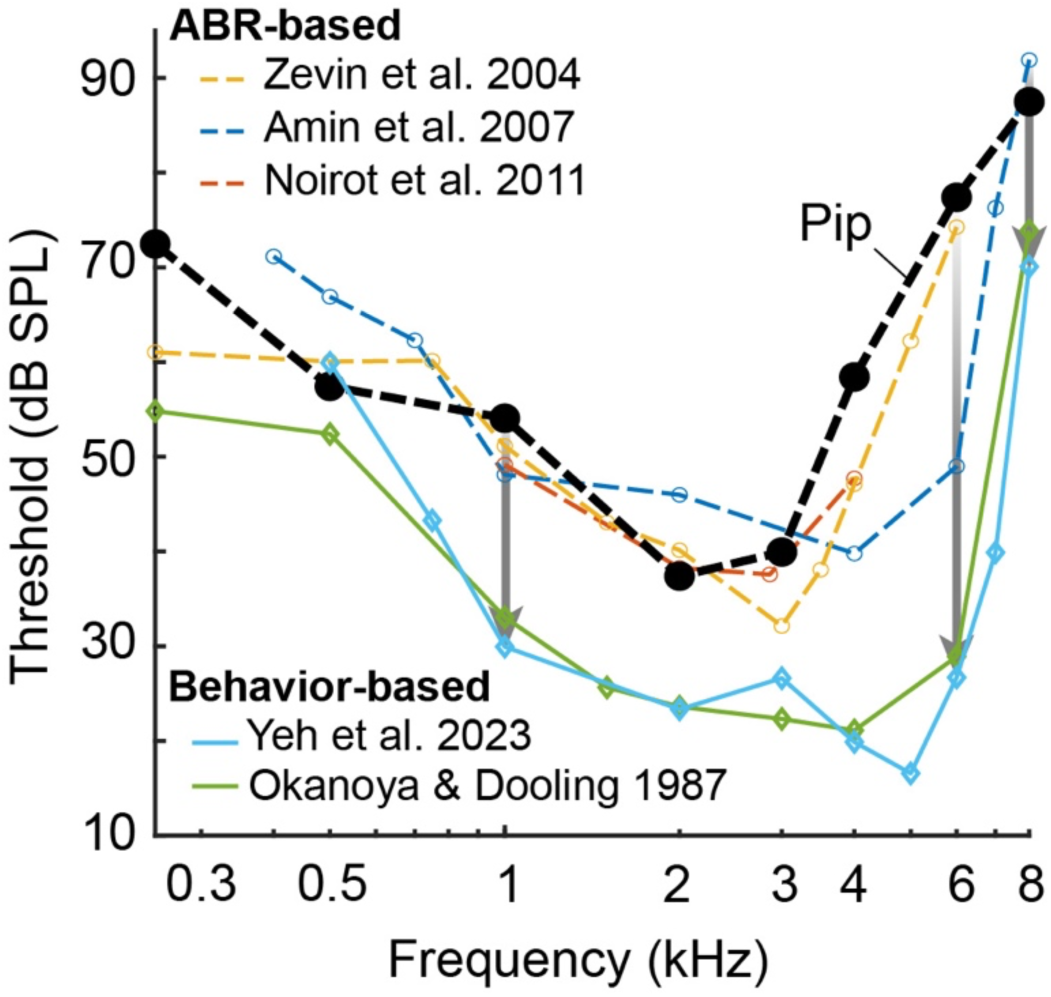
ABR and behavior-based audiograms in adult zebra finches are consistent in sensitivity and shape. The ABR based audiograms from four different laboratories including our pip-data are consistent in overall sensitivity and shape. The distinct low threshold at 6 kHz in the data by Amin et al. (2007) may reflect real biological variation between laboratory populations. The behavioral-based audiograms (Okanoya and Dooling, 1987; Yeh et al., 2023) are also consistent in shape and thresholds. The systematically lower threshold of the behavior-based audiograms varies from about 20 dB (downward arrows) in the 0.25 – 4 kHz range to 20-40 dB at 6 kHz.

Tone-burst thresholds in our dataset are higher than click-evoked thresholds, consistent with reduced neural synchrony for longer, frequency-specific stimuli, particularly at higher frequencies. The sensitivity in responses between tone pips and burst may be caused by different rise times. Our 5 ms tone pip stimuli had rise-fall times of 1 ms based on procedures in earlier studies (e.g. Brittan-Powell and Dooling, 2004; Ingham 2019). We used rise-fall times of 5 ms in our 25ms long tone bursts, because initial trials showed that 1 ms rise time resulted in click artefacts at low frequencies, where the rise-fall time is less than a cycle. Because initial measurements contained onset and offset responses, we increased the rise-fall times. Despite this limitation, these data provide a reliable within-method comparison across development. Sensitivity increases monotonically with age and emerges first at low frequencies, progressing toward higher frequencies. This functional trajectory matches patterns described across vertebrates (Saunders et al., 1973; Lippe, 1994; Jones et al., 2006). Hovewer it contrasts with an earlier ABR-based audiogram in zebrafnnches (Amin et al., 2007) and with the base-to-apex (high-to-low frequency) gradient of cochlear differentiation. This dissociation indicates that maturation of synaptic transmission, neural innervation, and central processing—rather than hair cell differentiation alone—limits early auditory function (Cohen and Fermin, 1978; Caus Capdevila et al., 2021). Future work using masked ABR may better resolve early frequency tuning (Brandt et al,. 2018).

ABR-derived audiograms systematically overestimate auditory thresholds relative to behavioral measures, because they require synchronous activation of large neural populations, whereas behavioral detection can rely on the most sensitive auditory nerve fibers (Pickles, 2012). Across taxa, this offset is typically on the order of 20 dB, depending on stimulus parameters and threshold criteria (Stapells, 2000; Brittan-Powell et al., 2002). This relationship is also observed in birds, including zebra finches, where behavioral thresholds are consistently lower than ABR thresholds but show similar overall tuning and frequency dependence (**Fig 4A**). Importantly, this offset does not alter the developmental trajectory we report: even accounting for a 20-40 dB difference, early hatchlings remain far less sensitive than adults, particularly at higher frequencies. Thus, while ABR underestimates absolute sensitivity, it provides a robust and conservative measure of functional auditory maturation across development.

The postnatal maturation of hearing may also affect the onset and efficacy of vocal imitation learning. In zebra finches, sensory song learning starts on 25 DPH (Immelmann, 1967; Böhner,1990; Roper and Zann, 2006; Braaten, 2010; Gobes et al., 2019; Leitão and Gahr, 2024). While the auditory sensitivity to click stimuli appears be mature at 10 DPH, the response amplitude of the auditory nerve (ABR wave I) increases past 20 DPH. Furthermore, ABR features that are generated by more upstream neural populations, e.g. wave II and later, appear to go through major changes after 10 DPH (**Fig 1C**). These data suggest that both peripheral auditory nerve and downstream circuits of the zebra finch auditory system continue to develop for several weeks after hatching. In chickens, the temporal coding ability of the auditory nuclei - the ability to respond to high stimulus rates – also increases post-hatch (Saunders et al., 1973, 1974; Manley et al., 1991; Warchol and Dallos, 1990). However, we know surprisingly little about these changes in zebra finches in early development (Hong and Sanchez, 2018) but they seem to continue well into the sensorimotor phase of song learning. The striking co-occurrence of the onset of the sensory phase of song learning with the maturation of the auditory nerve responses suggests that the onset of song learning in zebra finches may be gated by the developmental maturation of the auditory system. Furthermore, nascent, internal song templates acquired prior to the full maturation of hearing function may also require updating to match the changing target.

## Methods and Materials

### Animals

Local, captivity-bred zebra finches (*Taeniopygia guttata,* order *Passeriformes*) were used in this study. Zebra finches were kept and bred indoors in temperature- and humidity-controlled group aviaries (100-to-200 individuals) or pairwise in breeding cages supplemented with nesting boxes. Lights in the aviaries were on a 12/12 hour light/dark photoperiod. Zebra finches were given water, food, cuttlefish bones, and nesting material *ad libitum*. Nesting boxes were monitored daily for accurate aging of the hatched chicks. The day when a given zebra finch chick was observed to have hatched in the morning was defined to be post-hatch day 0 (0 DPH). Hatched chicks were recognized from each other by making individualized, non-invasive down feather cuts until the usage of leg bands was feasible at DPH 10. Weights for each age group are listed in **Table S6**. Animal experiments were approved by the Danish National Animal Experimentation Board (Dyreforsøgstilsynet) under license number 2019-15-0201-00284.

### Anesthesia

#### Preparation

30-to-60 minutes prior to recording, 20 DPH-to-adult zebra finches were removed from their breeding cage or aviary, brought to the test room in a transport cage, and were fasted to minimize the reflux of fluid/food during anesthesia. Younger zebra finches were moved to the test room in a warmed transport box and were not fasted. Zebra finches were weighed prior to the induction of anesthesia.

#### Dosing

Anesthesia was induced with an intrapectoral injection of ketamine (Ketaminol Vet., 100 mg/ml, MSD Animal Health, NJ, USA) in combination with xylazine (Rompun vet., 20 mg/ml, Bayer Animal Health GmbH, Leverkusen, Germany). 20 DPH-to-adult zebra finches were administered 50 µg of ketamine and 10 µg of xylazine per gram of body weight. 2-to-10 DPH zebra finches were given 50 µg of ketamine and 5 µg of xylazine per gram of body weight. After one hour, a supplementary dose (75% of the induction dose) was administered. In some zebra finches, a clear liquid - possibly due to fluid reflux or due to increased production of oral secretions - was noticed in their mouths within 10-to-15 minutes from anesthesia induction. To avoid airways from being obstructed, a small piece of paper tissue was inserted to the side of the mouth to drain the liquid.

### Auditory brainstem responses (ABRs)

#### Recording room

All recordings were performed by placing the zebra finch inside an audiometric room (T-room, C-A TEGNÉR AB, Bromma, Sweden) equipped with a self-built vibration isolation table with a padded surface. The walls of the room were padded with sound absorbing wedges (Classic Wedge 30, EQ Acoustics, UK).

#### Maintenance of body temperature

The body temperature of the animals was maintained by placing a heating pad warmed by circulating water (TP702, Gaymar Instruments Inc., Orchard Park, NY, USA) underneath them and placing a small cloth on top of their body. Body temperature monitoring was performed with a J-type temperature probe connected to a thermocouple measurement device (NI USB-TC01, National Instruments). The tip of the probe was coated with Sylgard (184 Silicone Elastomer Kit, Dow Europe GMBH, Wiesbaden, Germany), covered with vaseline, and was placed withing the cloaca. With 2-to-10 DPH zebra finches where the cloaca was too small for probe insertion the probe was placed under the wing. With this method we could keep body temperatures ±0.5°C around the target temperature during an experiment. Ambient temperature was 22°C and body temperatures for each age group are listed in **Table S6**. Animals were thus all 1) kept at constant temperatures (±0.5°C), 2) very close to normal body temperature (39.6-40.0°C and 3) at the same temperature (max difference: 0.6°C). All animals including very young hatchlings, thus did not suffer hypothermia that can affect ABR sensitivity (Rossi and Britt 1984; Doyle and Fria 1985). Conditions were thus consistent across all age groups, and ABR responses are thus directly comparable within this experiment.

#### Head placement

The head of the zebra finch was slightly elevated by placing it on top of wax mound (Surgident Periphery Wax, Kulzer, IN, US). The ends of a wedge-shaped notch on the wax were slightly pressed together so that the wax held the lower mandible of the zebra finch in place. The auricular feathers covering the left external auditory meatus in mature zebra finches were cut with scissors to allow unobstructed stimulation of the auditory system. Prior to electrode placement, the appropriate depth of anaesthesia was confirmed by the lack of responsiveness to a toe-pinch.

#### Electrodes

Three subdermal needle electrodes were placed subcutaneously on the head of the animal in the following configuration (**Fig. 1A**): (1) Active electrode: Dorsal to the left auditory meatus, pointing towards to the top of the head. (2) Inverting electrode: On top of the head along the midline, pointing towards the bill; (3) Ground electrode: Dorsal to the right auditory meatus, pointing towards to the top of the head. Of note, to compare these recordings to studies where the positions of the active and inverting electrodes are the opposite one must invert the recorded signal. The impedances of the electrodes were measured (RA4LI, Tucker-Davis Technologies, FL, USA) to be below 1 kOhm in 2-to-10 DPH zebra finches and below 3 kOhm in 20 DPH-to-adult zebra finches. Because we gathered ABRs with subdermal needles, there was no local cooling due to surgical exposure as may have been the case in sides where electrodes were places directly into the brain (e.g. Amin et al., 2007, Zevin et al., 2004).

#### Recording devices

Recorded evoked potentials were passed from the electrodes to a headstage (RA4LI) in series with a preamplifier (RA4PA) and with a digital signal processor (RM2, all from Tucker-Davis Technologies, FL, USA). RM2 was controlled with custom-written software QuickABR (Brandt et al., 2018; Lauridsen et al. 2021). Recorded responses were amplified by 74 dB and digitized with a sampling rate of 25 kHz (16 bits).

#### Sound stimulation

Sound stimulation was generated in the RM2 controlled by QuickABR, amplified (Cambridge Audio, Azur 740A Integrated Amplifier, London, UK), and emitted from a loudespeaker (Wharfedale Diamond 220, Wharfedale Ltd., Huntingdon, UK). The loudspeaker was string suspended 25 cm away from the animal, directly facing the left auditory meatus. Sound reflections were minimized by placing the loudspeaker at a 45° angle relative to the closest wall of the record ing room. To calibrate the loudspeaker, a ½” microphone (G.R.A.S. microphone type 26AK, G.R.A.S., Holte, Denmark) was suspended at the position of the head of the animal and connected to a microphone amplifier (G.R.A.S. Power Module Type 12AA, Gain 0, Linear filtering, Direct mode select = Ampl.). The microphone itself was calibrated using an acoustical calibrator (type 4321, Output: 1kHz at 94 dB re. 20 µPa, Brüel & Kjær). Calibration of the loudspeaker was controlled by QuickABR.

#### Stimuli

ABRs were elicited with broadband clicks (20 µs) and 25 ms-long tonebursts at 0.25 kHz, 0.5 kHz, 1 kHz, 2 kHz, 3 kHz, 4 kHz, 6 kHz, and 8 kHz. To minimize spectral splatter caused by abrupt on/offsets of stimuli, the rise/fall time (5 ms, cos^2^-gated) of the tonebursts was selected to accommodate at least one full cycle of the stimulus with the lowest frequency. To further increase the frequency specificity of the stimulus, the plateau time (15 ms) was selected to allow at least 3 full cycles of the stimulus with the lowest frequency of 250 Hz. Using a constant duration across frequencies ensures comparable stimulus energy across frequencies.

Additionally, 5 ms-long tone pips (1 ms rise/fall time, cos^2^-gated) were collected from adult zebra finches. Stimuli were sampled at 25 kHz and at a repletion rate of 25 Hz (clicks) and 3Hz (pips and bursts) to collect average responses consisting of 400 individual responses. ABR signal-to-noise ratio improves with the square root of the number of averages. Increasing from 400 to 1000 sweeps would reduce thresholds by at most ∼4 dB. Every second presented individual stimulus was inverted to reduce microphonic effects. The maximal sound pressure level (SPL) used in this study was selected to be 95 dB SPL as it can be viewed as being physiologically relevant. Click amplitudes are stated as the root mean square (RMS) dB value of a sinusoid with the same peak amplitude as the click (peak equivalent dB).

#### Recording paradigm

The stability of the recorded responses was monitored throughout the recording session by recording a 95 dB SPL click-evoked ABR periodically between each stimulus type. A specific stimulus recording was accepted into further analysis if the amplitude of the first click-evoked ABR wave had changed less than 25% from the most temporally stable amplitude of that recording session. To minimize test order bias, the recording order of stimuli with varying frequencies was randomized. Three animals were killed with an overdose of ketamine at the end of the recording session to verify the lack of response potentials *post mortem*.

#### ABR recovery from paired clicks

If consecutive clicks are played back at a too high rate, this can suppress the ABR response (Brittan-Powell and Dooling, 2004). To test the effects of stimulus repetition rate on the ABR, five adult zebra finches were presented at 6 Hz with paired-click and single click stimuli (Henry et al., 2021). The time between the first and second click (inter-click interval) was varied at 1 ms, 2 ms, 4 ms, 8 ms, 16 ms, 32 ms, and 48 ms. Each ABR was averaged from a total of 512 stimulus representations. The derived ABR to the second click was obtained by point-by-point subtraction of the single click stimulus ABR from the paired click stimulus ABR. To calculate the percentage of ABR recovery, the amplitude of the ABR wave I of the derived, second response was divided by the amplitude of the ABR wave I to the single click stimulus. These data show that the click rate of 25 Hz we used did not affect the magnitude of the ABR response (**Figure S3**). The burst rate of 3 Hz is a low presentation rate and is not expected to affect the ABR amplitude.

### Egg vibration measurements with laser doppler vibrometry

Zebra finch eggs (N = 5) were removed from their nest and placed in an upright position on a dental wax covered platform at the centre of an anechoic room. The anechoic room was custom-made with its walls and floor covered with 30 cm rockwool wedges. The sample platform was mounted on a heavy tripod which also held a laser Doppler vibrometer (OFV-5000 with OFV-505 sensor, Polytec, Waldbronn, Germany) attached on a manipulator. The laser was pointed at the centre of the side of the egg. Strong reflections were directly obtained from the egg surface so adding reflectors was not necessary. Sound-induced vibrations of the egg were induced with a speaker (JBL IG, Northridge, CA, USA) placed next to the laser vibrometer. A microphone (½ inch G.R.A.S., Brüel & Kjær) was placed few millimeters above the egg and was used for speaker calibration. The egg was stimulated with sound frequency sweeps (0.25-to-10 kHz) at 84 dB SPL. The sweeps were generated with Tucker-Davis system 2 hardware (Tucker-Davis Technologies, TDT, Alachua, FL, USA) controlled by custom-written software DragonQuest (Christensen-Dalsgaard and Manley, 2008).

The signal was deconvoluted by dividing the spectrum of the sweep with the measured transfer function of the speaker. The analog laser signal of the laser vibrometer was digitized using an A–D converter (AD2, TDT) and averaged over 100 presentations. During recordings, the laser signal was monitored using an oscilloscope and a vibrometer controller (Polytec OFV-5000). Stimulation and recording were controlled by DragonQuest.

### Data analysis and visualization

Data was visualized and analysed with custom scripts in MATLAB R2022B (Mathworks). Figures were edited with Illustrator CS6 (Adobe).

#### Threshold determination

For ABR threshold detection, the stimulus level was lowered from the maximal value in 5 dB steps. For the reported thresholds to reflect the average sensitivity of zebra finches capable of detecting the given stimulus, only zebra finches that displayed a detectable ABR (both visually and above the signal-to-noise ratio of 2) at the maximal stimulation level are included in the analysis. The response threshold was designated to be 2.5 dB below the last observable response-generating stimulus level (*arrow* in **Fig 1F**).

#### Signal-to-noise ratio calculation

The recording prior to stimulus onset was used to calculate the noise RMS. Signal-to-noise ratio (S/N) was determined by dividing the RMS value of the signal (*blue window*) by the RMS of the recording prior to stimulus onset (*red window*, **Figs. 1D; 2A,B** and **Fig. S2A**). We used an automated criteria detection of thresholds to remove potential experimenter bias and checked the automatic thresholds by visual inspection.

#### Response latency and amplitude calculation

Response latency was calculated as the time difference between the first negative deflection of the ABR (designated as wave I) and the arrival time of the sound at the ear (**Fig 1C**). The amplitude of the wave I (reflecting the activity of the auditory nerve) was calculated as the potential difference between the first negative peak and the following positive trough that together formed the most prominent component of the response (**Fig 1C**). Of note, a second negative peak/deflection between the first peak and the trough existed. This peak appeared to merge with the first negative peak at soft SPLs as reported for ABRs of bald eagles and red-tailed hawks (McGee et al., 2019).

### Statistics

To test for main effects, we used linear mixed effects models. We sequentially compared models by adding hypothesised effects as fixed effects one-by-one, starting from a null model and ending with full interaction models. For all age comparisons, age was treated as an ordinal rather than a continuous variable to account for discrete timepoints of measurements and to allow for post-hoc pairwise comparisons between ages. All models had the individual animals as random effect. We used chi-square tests, Akaike Information Criterion (AIC), and Bayes Information Criterion (BIC) as criteria for model comparison. For models with significant main effects, we then used estimated marginal means for post-hoc comparisons. We used Bonferroni p-value correction to account for multiple comparisons. In all figures, we denote significance levels as follows: *, p(Bonf.) = 0.05; **, p(Bonf.) = 0.01; ***, p(Bonf.) = 0.001. All statistical testing was carried out in R 4.2.1 (R Core Team, 2024) We used the lme4 package (Bates et al., 2015) for mixed effects modelling and emmeans package (Lenth, 2024) for post-hoc comparisons.

## Acknowledgments

We thank Catherine Carr and Albertine Leitão for their feedback on the manuscript. We also thank Sonja Jacobsen, Emilie Radoor, Emilie Jensen, Dina Stær Arengoth, and Niels Peter Jørgensen for animal care and maintenance.

## Author contributions

T.A., J.C.D., and C.P.H.E. conceptualized experiments; T.A. collected and analyzed the data for figures 1, 2, S1, S2 and S3; T.A. and J.C.D. collected and analyzed the data for figure 3; T.A. and C.P.H.E. wrote the draft of the manuscript; T.A., J.C.D., and C.P.H.E. wrote the manuscript; T.A. and C.P.H.E. acquired funding.

