## Supplemental Information for "Progressive postnatal hearing development limits early parent-offspring vocal communication in the zebra finch"

### Supplemental figures

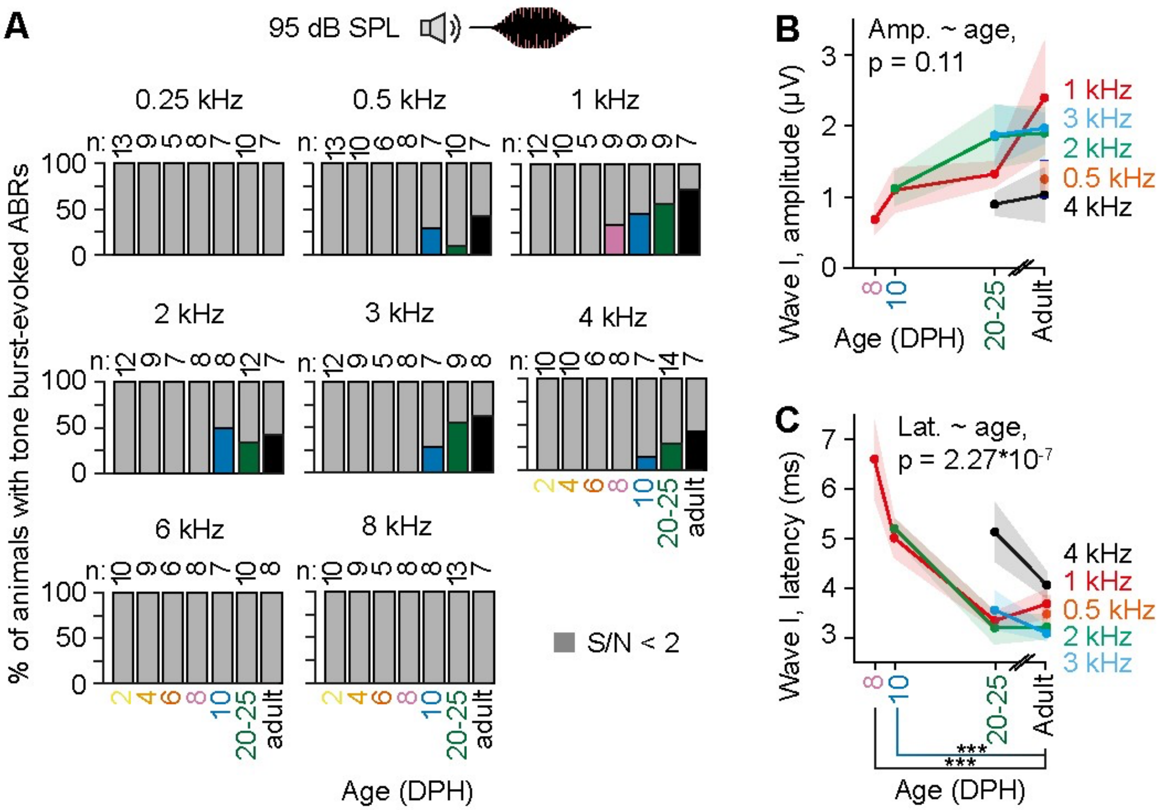


**Figure S1. Development of tone burst-evoked auditory brainstem responses.** (**A**) Percentage of animals displaying tone burst-evoked ABRs at 95 dB SPL. Gray bars indicate animals with S/N < 2. Exact number of animals is displayed above the bars. (**B**) Changes in tone burst-evoked ABR wave I amplitude and (**C**) latency over postnatal development. Frequencies are color coded as displayed below the figure. Shaded areas in B,C = s.e.m.

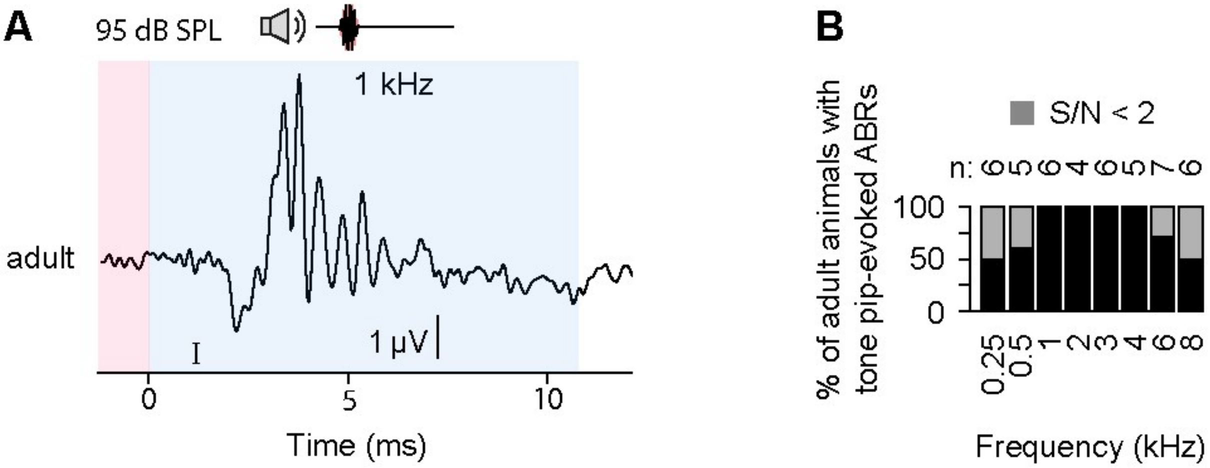


**Figure S2. Tone pip-evoked auditory brain stem responses in adult zebra finches** (**A**) Representative AC-ABR evoked by a 95 dB SPL, 1 kHz tone pip recorded from an adult zebra finch. (**B**) Percentage of adult zebra finches showing tone burst-evoked ABRs at 95 dB SPL between 0.25 and 8 kHz. Gray bars indicate responses with S/N < 2.

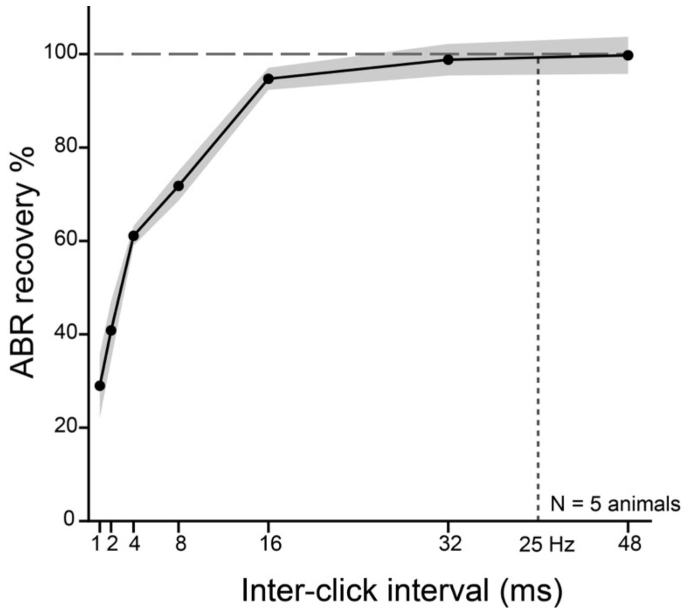


**Figure S3. ABR response is not suppressed by inter click intervals above 32 ms.**

### Supplemental tables

**Table S1. Statistics for click-evoked auditory brainstem response thresholds**

| Click threshold: Model comparisons (Fig. 1g) | | | | | |
|---|---|---|---|---|---|
| **Model** | **Residual degrees of freedom** | **Residual sum of squares** | **Degrees of freedom** | **Sum of squares** | **F value** |
| Null model | 60 | 17345.190 | | | |
| Age | 54 | 4561.668 | 6 | 12783.523 | 25.221 |

| Click threshold: Coefficients (Fig. 1g) | | | | |
|---|---|---|---|---|
| **Parameter** | **Parameter estimate** | **Standard error** | **t value** | **p value (t)** |
| Intercept | 54.592 | 1.204 | 45.329 | 1.162E-44 |
| Age | -37.327 | 3.273 | -11.405 | 5.320E-16 |

| Click threshold: Post-hoc comparisons (Fig. 1g) | | | | | |
|---|---|---|---|---|---|
| **Contrast** | **Parameter estimate** | **Standard error** | **Degrees of freedom** | **t ratio** | **p value (t)** |
| 4 DPH vs. adult | 37.500 | 4.596 | 54 | 8.160 | 3.268E-10 |
| 6 DPH vs. adult | 29.375 | 4.964 | 54 | 5.918 | 1.385E-06 |
| 8 DPH vs. adult | 14.375 | 4.271 | 54 | 3.366 | 0.008 |
| 10 DPH vs. adult | 5.284 | 4.271 | 54 | 1.237 | 1 |
| 20 DPH vs. adult | -0.639 | 4.757 | 54 | -0.134 | 1 |
| 25 DPH vs. adult | -5.625 | 4.360 | 54 | -1.290 | 1 |

**Table S2. Statistics for click-evoked auditory brainstem response amplitudes**

**Click amplitude: Model comparisons (Fig. 1h)**

| Model | Number of parameters | AIC | BIC | Log-likelihood | Deviance | Chi square | Degrees of freedom | p value (chi square) |
|---|---|---|---|---|---|---|---|---|
| Null model | 3 | 2712.939 | 2725.723 | -1353.469 | 2706.939 | | | |
| Fixed effect: SPL | 4 | 2276.820 | 2293.866 | -1134.410 | 2268.820 | 438.119 | 1 | 2.779 E-97 |
| Fixed effects: SPL + Age | 10 | 2209.391 | 2252.006 | -1094.695 | 2189.391 | 79.429 | 6 | 4.687 E-15 |
| Interaction SPL * Age | 16 | 2057.162 | 2125.345 | -1012.581 | 2025.162 | 164.229 | 6 | 7.524 E-33 |

**Click amplitude: Coefficients (Fig. 1h)**

| Parameter | Estimate | Standard error | Degrees of freedom | t value | p value (t) |
|---|---|---|---|---|---|
| Intercept | -4.536 | 0.800 | 508.707 | -5.670 | 2.402E-08 |
| SPL | 0.106 | 0.009 | 480.584 | 11.972 | 4.339E-29 |
| Age | -5.626 | 2.796 | 508.128 | -2.012 | 4.472E-02 |
| SPL * Age | 0.166 | 0.031 | 481.530 | 5.356 | 1.323E-07 |

**Click amplitude: Post-hoc comparisons (Fig. 1h)**

| Contrast | Estimate | Standard error | Degrees of freedom | t ratio | p value (t) |
|---|---|---|---|---|---|
| 4 DPH vs. adult | -6.750 | 1.250 | 154.026 | -5.398 | 1.498E-06 |
| 6 DPH vs. adult | -6.109 | 1.082 | 70.689 | -5.649 | 1.891E-06 |
| 8 DPH vs. adult | -5.409 | 0.859 | 52.494 | -6.296 | 3.785E-07 |
| 10 DPH vs. adult | -4.723 | 0.854 | 51.324 | -5.530 | 6.533E-06 |
| 20 DPH vs. adult | -2.850 | 0.951 | 51.361 | -2.995 | 0.025 |
| 25 DPH vs. adult | -0.601 | 0.871 | 51.122 | -0.690 | 1 |

**Table S3. Statistics for click-evoked auditory brainstem response latencies**

**Click latency: Model comparisons (Fig. 1i)**

| Model | Number of parameters | AIC | BIC | Log-likelihood | Deviance | Chi square | Degrees of freedom | p value (chi square) |
|---|---|---|---|---|---|---|---|---|
| Null model | 3 | 833.089 | 845.810 | -413.544 | 827.089 | | | |
| Fixed effect: SPL | 4 | 80.851 | 97.812 | -36.425 | 72.851 | 754.238 | 1 | 4.808 E-166 |
| Fixed effects: SPL + age | 10 | -16.518 | 25.884 | 18.259 | -36.518 | 109.369 | 6 | 2.763 E-21 |
| Interaction: SPL * age | 16 | -120.212 | -52.367 | 76.106 | -152.212 | 115.693 | 6 | 1.306 E-22 |

**Click latency: Coefficients (Fig. 1i)**

| Parameter | Estimate | Standard error | Degrees of freedom | t value | p value (t) |
|---|---|---|---|---|---|
| Intercept | 6.911 | 0.108 | 365.521 | 64.005 | 1.111 E-200 |
| SPL | -0.030 | 0.001 | 451.876 | -28.302 | 4.037 E-102 |
| Age | -4.290 | 0.359 | 453.952 | -11.964 | 7.319E-29 |
| SPL * Age | 0.021 | 0.004 | 452.408 | 5.809 | 1.185E-08 |

**Click latency: Post-hoc comparisons (Fig. 1i)**

| Contrast | Estimate | Standard error | Degrees of freedom | t ratio | p value (t) |
|---|---|---|---|---|---|
| 4 DPH vs. adult | 3.080 | 0.245 | 73.488 | 12.560 | 3.405 E-19 |
| 6 DPH vs. adult | 2.495 | 0.248 | 57.938 | 10.063 | 1.491 E-13 |
| 8 DPH vs. adult | 1.522 | 0.210 | 52.527 | 7.242 | 1.159 E-08 |
| 10 DPH vs. adult | 1.297 | 0.210 | 52.263 | 6.178 | 5.945 E-07 |
| 20 DPH vs. adult | 0.520 | 0.232 | 52.279 | 2.239 | 0.177 |
| 25 DPH vs. adult | 0.357 | 0.214 | 52.221 | 1.668 | 0.608 |

**Table S4. Statistics for tone burst-evoked auditory brainstem response amplitudes**

**Tone burst amplitude: Model comparisons (Fig. 2a and Supplementary Fig. S1b)**

| Model | Number of parameters | AIC | BIC | Log-likelihood | Deviance | Chi square | Degrees of freedom | p value (chi square) |
|---|---|---|---|---|---|---|---|---|
| Null model | 3 | 117.202 | 122.688 | -55.601 | 111.202 | | | |
| Fixed effect: Age | 6 | 117.095 | 128.067 | -52.547 | 105.095 | 6.107 | 3 | 0.107 |

**Tone burst latency: Model comparisons (Fig. 2a and Supplementary Fig. S1c)**

| Model | Number of parameters | AIC | BIC | Log-likelihood | Deviance | Chi square | Degrees of freedom | p value (chi square) |
|---|---|---|---|---|---|---|---|---|
| Null model | 3 | 150.112 | 155.598 | -72.056 | 144.112 | | | |
| Fixed effect: Age | 6 | 122.392 | 133.364 | -55.196 | 110.392 | 33.720 | 3 | 2.270E-07 |

**Tone burst latency: Coefficients (Fig. 2a and Supplementary Fig. S1c)**

| Parameter | Estimate | Standard error | Degrees of freedom | t value | p value (t) |
|---|---|---|---|---|---|
| Intercept | 4.744 | 0.159 | 42 | 29.826 | 6.790E-30 |
| Age | -2.392 | 0.361 | 42 | -6.631 | 4.920E-08 |

**Tone burst latency: Post-hoc comparisons (Fig. 2a and Supplementary Fig. S1c)**

| Contrast | Estimate | Standard error | Degrees of freedom | t ratio | p value (t) |
|---|---|---|---|---|---|
| 8 DPH vs. adult | 3.133 | 0.527 | 38.725 | 5.951 | 1.866E-06 |
| 10 DPH vs. adult | 1.640 | 0.363 | 25.072 | 4.522 | 0.0004 |
| 20-to-25 DPH vs. adult | 0.342 | 0.294 | 10.697 | 1.161 | 0.813 |

**Tone burst threshold: Model comparisons (Fig. 2c)**

| Model | Number of parameters | AIC | BIC | Log-likelihood | Deviance | Chi square | Degrees of freedom | p value (chi square) |
|---|---|---|---|---|---|---|---|---|
| Null model | 3 | 353.752 | 359.238 | -173.876 | 347.752 | | | |
| Fixed effect: Age | 6 | 358.107 | 369.079 | -173.054 | 346.107 | 1.645 | 3 | 0.649 |

**Table S5.** Statistics for comparing tone pip- vs tone burst-induced auditory brainstem responses

| Tone pip vs tone burst: Model comparisons (Fig. 2c and Supplementary Fig. 2) | | | | | | | | |
|---|---|---|---|---|---|---|---|---|
| **Model** | **Number of parameters** | **AIC** | **BIC** | **Log-likelihood** | **Deviance** | **Chi square** | **Degrees of freedom** | **p value (chi square)** |
| Null model | 3 | 477.716 | 483.683 | -235.858 | 471.716 | | | |
| Fixed effect: Stimulus type | 4 | 469.444 | 477.400 | -230.722 | 461.444 | 10.272 | 1 | 0.00135 |
| Fixed effects: Stimulus type + Frequency | 5 | 464.609 | 474.554 | -227.305 | 454.609 | 6.835 | 1 | 0.00894 |

| Tone pip vs tone burst: Coefficients (Fig. 2c and Supplementary Fig. 2) | | | | | |
|---|---|---|---|---|---|
| **Parameter** | **Estimate** | **Standard error** | **Degrees of freedom** | **t value** | **p value (t)** |
| Intercept | 68.849 | 4.706 | 26.832 | 14.631 | 2.628E-14 |
| Stimulus type | -19.651 | 4.878 | 50.590 | -4.028 | 0.000189 |
| Frequency | 0.00311 | 0.00111 | 50.027 | 2.786 | 0.00751 |

**Table S6. Weights and temperature during ABR measurements for all age groups.** Abbreviations: DPH, Days post-hatch; SD, Standard deviation; SEM, Standard error of the mean; N, number of animals.

***Weight of tested animals***

| | DPH 2 | DPH 4 | DPH 6 | DPH 8 | DPH 10 | DPH 20 | DPH 25 | Adult |
|---|---|---|---|---|---|---|---|---|
| **Mean weight:** | 1.6 | 3.1 | 4.2 | 7.0 | 10.1 | 13.5 | 14.0 | 15.8 |
| **SD:** | 0.4 | 0.8 | 1.3 | 1.3 | 1.5 | 1.0 | 1.3 | 1.4 |
| **SEM:** | 0.1 | 0.2 | 0.4 | 0.4 | 0.5 | 0.4 | 0.4 | 0.5 |
| **N:** | 14 | 13 | 10 | 11 | 11 | 7 | 8 | 8 |

***Temperature of tested animals***

| | DPH 2 | DPH 4 | DPH 6 | DPH 8 | DPH 10 | DPH 20 | DPH 25 | Adult |
|---|---|---|---|---|---|---|---|---|
| **Mean temperature:** | 39.5 | 39.6 | 40.0 | 39.7 | 39.6 | 39.8 | 40.0 | 39.4 |
| **SD:** | 0.7 | 0.6 | 0.7 | 0.7 | 0.8 | 0.7 | 1.3 | 1.3 |
| **SEM:** | 0.2 | 0.2 | 0.2 | 0.2 | 0.2 | 0.3 | 0.5 | 0.5 |
| **N:** | 14 | 13 | 10 | 11 | 11 | 7 | 8 | 8 |
